# Compartment-dependent selection regimes maintain the G-quadruplex paradox across the eukaryotic kingdoms

**DOI:** 10.64898/2026.05.21.727063

**Authors:** Masato Tanigawa, Takafumi Iwaki

## Abstract

G-quadruplex (G4) DNA structures are depleted from coding sequences (CDS) and enriched in regulatory regions across diverse genomes (the “G4 paradox”), but the selection regime maintaining this architecture across eukaryotes remains unclear. Analysing 198 reference genomes balanced across the four kingdoms (Animalia 86, Fungi 49, Plantae 33, Protozoa 30), we propose a *two-regime architecture* in which the dominant selective force depends on the genomic compartment. In CDS, codon-adaptation selection takes priority: naive Nei–Gojobori comparison shows 5–16% synonymous-rate (*d_S_*) suppression at G4 codons in six of seven unsaturated species pairs, but the signal is absorbed by per-gene codon-adaptation-index (CAI) adjustment (0*/*7 pairs retain a G4 odds ratio below 1; five reverse to OR *>* 1). In introns and promoters, where no codons exist, selection acts directly on G4 structure: phylogenetic generalised least-squares (PGLS) yields five Benjamini–Hochberg-significant kingdom-specific helicase associations (*q <* 0.10), all at intronic or promoter G4s, including context-dependent FANCJ/BRIP1 (*β* = +1.60, *q* = 0.057 for intron G4 in Plantae; *β* = *−*2.49, *q* = 0.004 for promoter G4 in Protozoa); three of five remain significant on an independent Open Tree of Life backbone. Intronic G4 are concentrated in deeply conserved orthologue groups (*≥* 87% in every kingdom), consistent with maintenance of regulatory G4 host-gene context since early eukaryotic evolution.

**Interim-correction notice (bioRxiv v3, 2026-08-03):** This preprint version corrects a set of documentation-level errors identified in a coauthor audit of the analysis pipeline (available as AUDIT_REPORT.md in the Zenodo deposit). Several central analytical claims of this manuscript — the definition of the log_2_ IRR (density-vs. fraction-ratio), the downstream use of consensus-stable versus candidate G4 motifs, the CAI-controlled *d_S_* adjustment model, the filter chain in the stable-G4 classifier, and the reconciliation of PGLS output across main, alternative-tree, robustness, and Supplementary Information files — are currently under clean-environment re-verification. The outcome of that re-verification may change specific numerical values, the direction of some regional sign summaries, and the framing of downstream conclusions in a subsequent version. Readers should treat the present version as an interim documentation correction, not as a fully verified revision. Users planning to build on the deposited results should consult CORRECTION_NOTES.md and AUDIT_REPORT.md in the Zenodo deposit for the current list of open methodology-level issues.

## Introduction

G-quadruplex (G4) DNA structures are non-canonical four-stranded conformations formed by guanine-rich sequences via Hoogsteen base-pairing of four guanines stacked around a central potassium ion (Sen and Gilbert, 1988; Burge et al., 2006). They occupy a singular position in molecular biology: kinetically stable G4 within coding sequences can stall replication forks and RNA polymerase, generating mutational and transcriptional liabilities, yet the same structures are repeatedly enriched in promoters, untranslated regions, and immunoglobulin loci across distantly related eukaryotes, where their stability appears to be exploited rather than purged (Maizels and Gray, 2013; Hänsel-Hertsch et al., 2017, 2018; Spiegel et al., 2020).

This dual character has been progressively documented across the tree of life. Cross-species surveys have reported preferential positioning of G4 motifs in regulatory regions (Huppert and Balasubramanian, 2007; Eddy and Maizels, 2008); the Quadrupia catalogue has extended this signature to 108,449 genomes (Chantzi et al., 2025); and Puig Lombardi et al. (2019) demonstrated systematic depletion of thermodynamically stable G4 motifs across 600 species, implying purifying selection against deleterious G4 in coding sequences. Our recent analysis of 31 coronavirus genomes (Tanigawa and Iwaki, 2026) described the same tension as a *G4 paradox*: simultaneous whole-genome G4 depletion under a dinucleotide-preserving null and pronounced enrichment within specific functional regions of the same genome. The present manuscript provides a rigorous, kingdom-balanced cross-eukaryotic test of that observation.

Three questions structure the present analysis. First, is the G4 paradox a genuine cross-kingdom property of eukaryotic genome architecture, or an Animalia-biased artefact of historical sampling effort? Second, what molecular machinery (G4-resolving helicases) co-evolves with the G4 burden across kingdoms, and does its deployment depend on the genomic compartment (CDS, intron, promoter) in which the G4 resides? Third, to what extent can putative codon-level mechanisms of CDS G4 maintenance, such as a structural constraint at silent sites, be inferred from comparative *d_N_ /d_S_* analysis once standard expression-coupled confounders are controlled?

We address all three by analysing a balanced sample of 198 reference genomes spanning the four eukaryotic kingdoms (Animalia 44%, Fungi 25%, Plantae 16%, Protozoa 15%), applying a hybrid G4 stability classifier (regex + G4Hunter consensus, Δ*G*_ML_ *≤ −*5 kcal/mol), performing Nei–Gojobori *d_N_ /d_S_* analysis at G4-overlapping codons versus control codons (both without and with per-gene codon-adaptation index control) for closely related species pairs within each kingdom, and quantifying G4 helicase orthologue counts (KEGG K10901 BLM/WRN/RECQ, K15364 FANCJ/BRIP1, K11271 RTEL1, K12823 DDX5) under phylogenetic correction (PGLS) on the curated species tree, with robustness checks on a polytomy-free subset and on an independently constructed Open Tree of Life synthetic backbone. The data confirm the G4 paradox as a cross-kingdom regulatory architecture sustained primarily by kingdom-specific helicase coevolution at intronic and promoter G4s, while the apparent CDS *d_S_* suppression at G4 codons is absorbed by CAI adjustment and does not constitute independent evidence for a standalone silent-site constraint.

## Results

### A balanced cross-kingdom sample confirms the G4 paradox as a sign-consistent genome feature across the four eukaryotic kingdoms

To rigorously test whether the G4 paradox is cross-kingdom across eukaryotes or biased by historical sampling effort, we assembled a balanced sample of 198 reference genomes spanning all four eukaryotic kingdoms: Animalia (*n* = 86, 43% of the sample, vs. 80% in Puig Lombardi et al., 2019), Fungi (*n* = 49, 25%), Plantae (*n* = 33, 16%), and Protozoa (*n* = 30, 15%) (Figure 2). Within Animalia, phylum-level diversity was enforced (Nematoda 31, Platyhelminthes 14, Arthropoda 23, Vertebrata 12, Annelida 6). Within Plantae, all major land-plant lineages were represented including a chromosome-scale *Ginkgo biloba* assembly (Ginkgoales, Gymnospermae; Liu et al., 2021) recon-structed from the Ginkgo Database (Gu et al., 2022) together with grasses, eudicots, bryophytes, lycophytes, and Charophyta.

For each species we extracted six genomic regions (5*^′^*UTR, CDS, intron, 3*^′^*UTR, promoter [*−*1000 to *−*1 relative to TSS], and intergenic) from a reference GFF, scanned them for G4 motifs with a regex–G4Hunter consensus classifier (Bedrat et al., 2016; Tanigawa and Iwaki, 2026), and stratified the calls by Mergny–Lacroix predicted thermodynamic stability (Δ*G ≤ −*5 kcal/mol for the *consensus stable* class; Mergny and Lacroix, 2003). The kingdom-stratified medians of the log_2_-enrichment ratio (log_2_ IRR; stable G4 density in each region versus the genome-wide background) are summarised in Table 1.

**Table 1:**
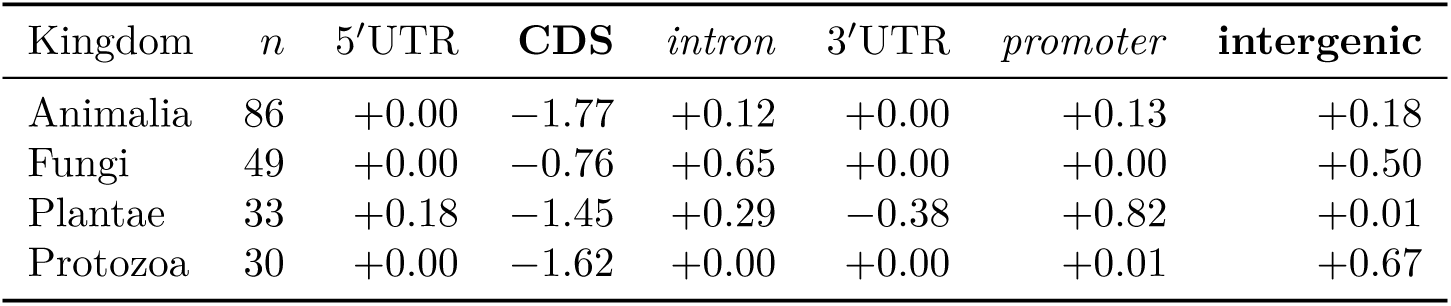
Median log_2_ IRR of consensus stable G4 motifs by genomic region and kingdom (final. *n* = 198). Positive values indicate enrichment versus the genome-wide background; negative values indicate depletion. Bold cells highlight the two regions (CDS, intergenic) whose kingdom medians are strictly sign-consistent across all four kingdoms (Rule B; Methods). Intron and promoter (italics) become sign-consistent under a tie-permissive rule (Rule A) that treats a kingdom median of zero as compatible with same-signed medians in other kingdoms; ties arise when a substantial fraction of species in the kingdom has zero stable G4 in that region (Protozoa intron: 15/30 species with zero stable G4; Fungi promoter: 8/49 with zero stable G4, and the kingdom median falls at zero because of ties around the background level).

| Kingdom | $n$ | 5'UTR | **CDS** | *intron* | 3'UTR | *promoter* | **intergenic** |
| --- | --- | --- | --- | --- | --- | --- | --- |
| Animalia | 86 | +0.00 | −1.77 | +0.12 | +0.00 | +0.13 | +0.18 |
| Fungi | 49 | +0.00 | −0.76 | +0.65 | +0.00 | +0.00 | +0.50 |
| Plantae | 33 | +0.18 | −1.45 | +0.29 | −0.38 | +0.82 | +0.01 |
| Protozoa | 30 | +0.00 | −1.62 | +0.00 | +0.00 | +0.01 | +0.67 |

### Two regions are strictly sign-consistent across all four kingdoms

Under the strict rule that all four kingdom medians agree in sign (Rule B; Methods), CDS depletion (range *−*0.76 to *−*1.77 log_2_ IRR) and intergenic enrichment (+0.01 to +0.67) are strictly sign-consistent (Figure 1); the joint probability of observing *≥* 2 of the six regions sign-consistent under a binomial null with per-region *P* = 0.0625 (one-sided, *n* = 4 kingdoms) is *P* = 0.049. Under a tie-permissive rule (Rule A) that treats a kingdom median of zero as compatible with same-signed medians in other kingdoms, intron and promoter enrichment become sign-concordant as well: intron (+0.00 to +0.65; Protozoa median 0.00) and promoter (+0.00 to +0.82; Fungi median 0.00) each show non-negative medians in all four kingdoms, yielding four sign-concordant regions with joint *P* = 2 *×* 10*^−^*^4^ (binomial, per-region *P* = 0.0625). The two rules therefore agree on a robust CDS-depletion / intergenic-enrichment pattern and disagree on the framing of intron and promoter enrichment, where the sign concordance depends on how kingdoms with median-at-zero (15/30 Protozoa species have zero stable intronic G4; 8/49 Fungi species have zero stable promoter G4, with the kingdom median at zero because of ties around the background level) are treated. We refer to CDS and intergenic as *robust cross-kingdom regions* and to intron and promoter as *tie-boundary cross-kingdom regions*; the two-tier pattern that emerges from either rule is robust to the sample rebalancing (Animalia reduced from 80% to 44% relative to v1) and is inconsistent with the hypothesis that the paradox is an Animalia-specific artefact.

**Figure 1:**
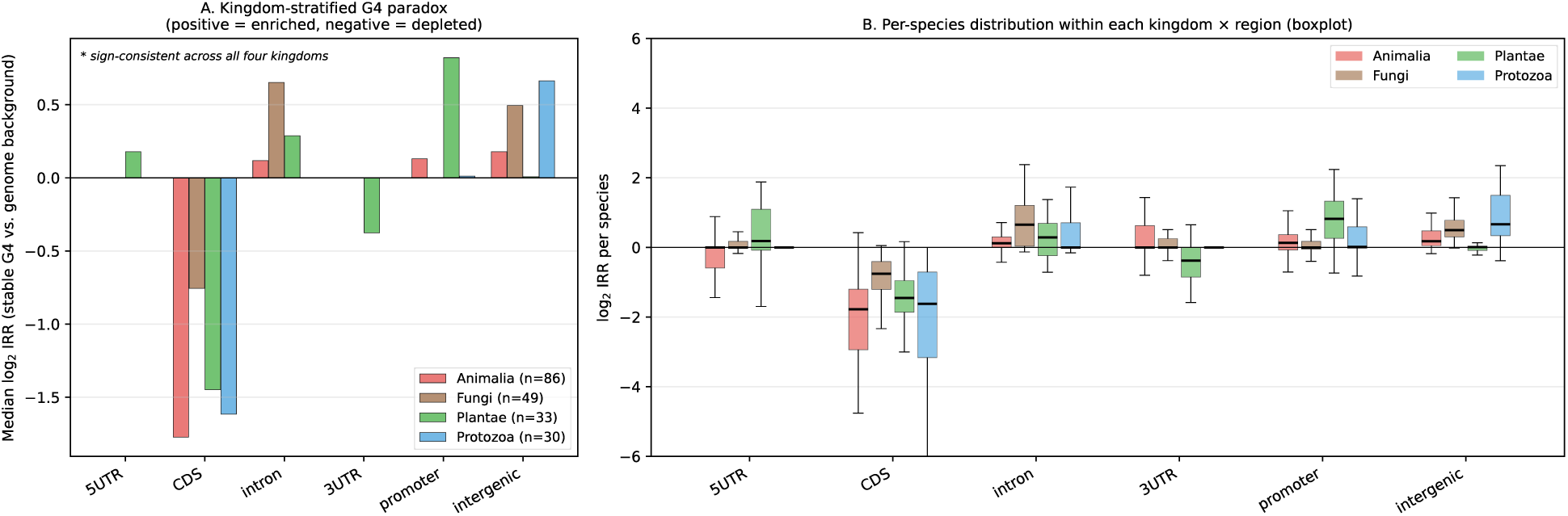
Cross-kingdom G-quadruplex paradox across 198 reference genomes. (A) Median log_2_ enrichment ratio (IRR) of consensus-stable G4 motifs per region, stratified by kingdom (Animalia *n* = 86, Fungi *n* = 49, Plantae *n* = 33, Protozoa *n* = 30). Asterisks mark the four regions whose kingdom medians share the same non-negative or non-positive sign under the tie-permissive Rule A (Methods): CDS depletion (strict; all four medians *<* 0), intergenic enrichment (strict; all four medians *>* 0), and intron and promoter enrichment (tie-permissive; three kingdom medians *>* 0 with one tie at exactly 0). Only CDS and intergenic satisfy the strict Rule B in which all four kingdom medians must be non-zero. (B) Per-species distribution (boxplot) of log_2_ IRR by region *×* kingdom, demonstrating that the kingdom-stratified medians in panel A reflect coherent within-kingdom trends rather than averaging artefacts.

**Figure 2:**
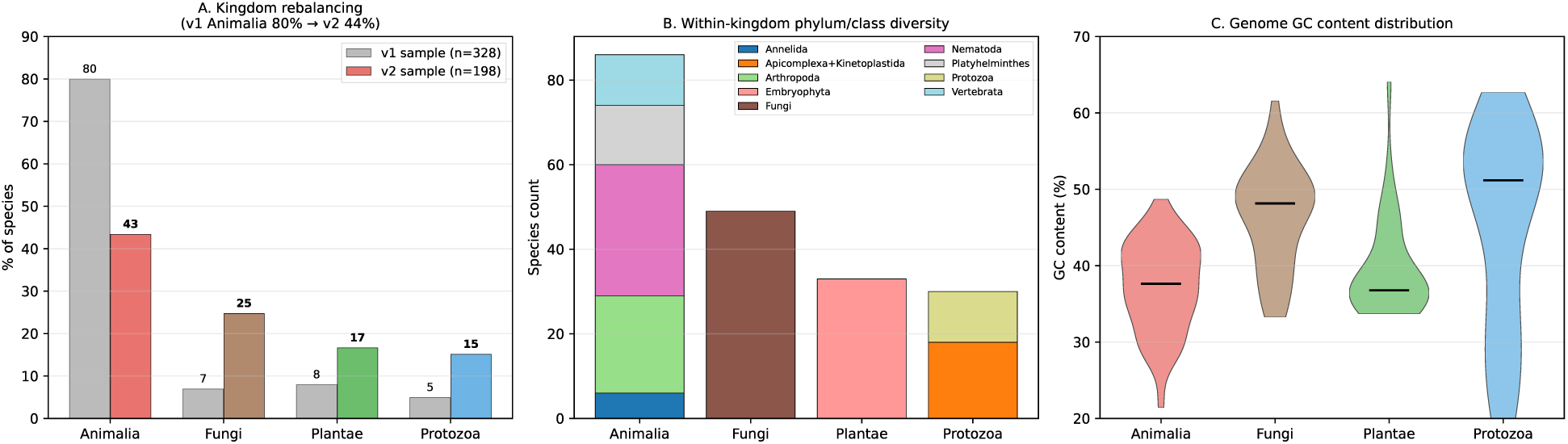
Composition of the rebalanced v2 sample. (A) Kingdom representation in v1 (*n* = 328, Animalia 80%) vs. v2 (*n* = 198, Animalia 43%), highlighting the systematic rebalancing. (B) Within-kingdom phylum/class diversity. Within Animalia, five phyla are represented (Nematoda, Platyhelminthes, Arthropoda, Vertebrata, Annelida); within Plantae, all major land-plant lineages are present together with charophyte and chlorophyte outgroups; Fungi and Protozoa span multiple major classes. (C) Genome GC content distribution per kingdom (violin plot); the four kingdoms span GC *∼*0.19–0.64, providing a broad GC range against which to test the GC dependence of G4 paradox metrics (Results).

**Figure 3:**
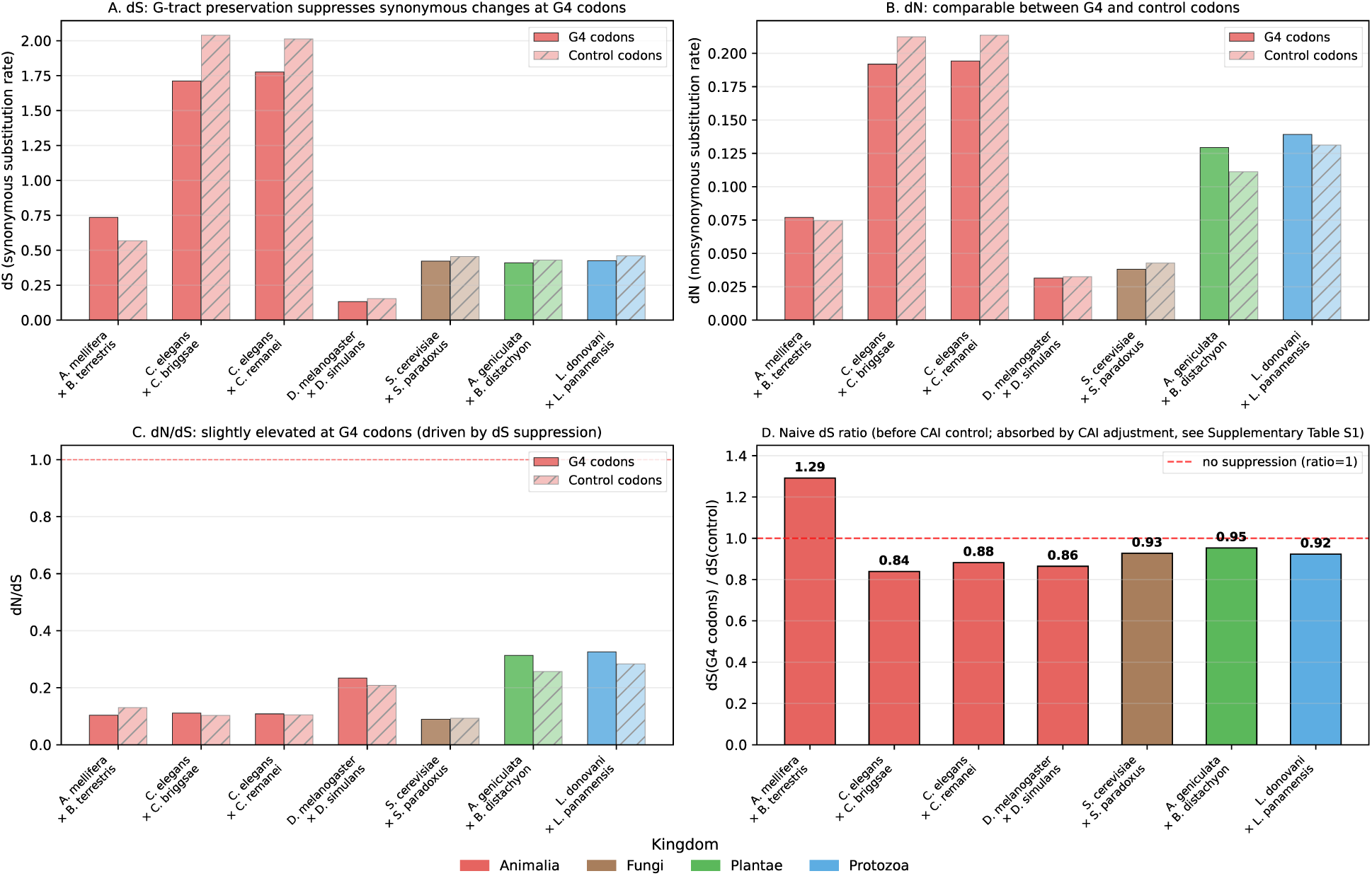
Naive Nei–Gojobori synonymous (*d_S_*) and nonsynonymous (*d_N_*) substitution rates at G4-overlapping versus control codons, before per-gene codon-adaptation-index (CAI) adjustment. (A) *d_S_* is lower at G4 codons than at control codons in 4/5 displayed pairs (ratio 0.84–0.95); the *Apis × Bombus* hymenopteran pair is the exception (1.29). (B) *d_N_* is comparable between G4 and control codons in all pairs. (C) *d_N_ /d_S_* at G4 codons is comparable to or slightly elevated over control. (D) *d_S_*(G4) / *d_S_*(control) ratio. These ratios faithfully describe the gene-pool synonymous rate at G4 codons; however, the CAI-controlled per-codon and per-gene analyses presented in the subsection below show that this apparent *d_S_* suppression is largely absorbed once per-gene CAI is included as a covariate, indicating that G4-overlapping codons preferentially reside in highly-expressed, codon-optimised genes. The values in panels (A) and (D) should therefore be read as a confounded readout rather than as a direct measure of structural silent-site constraint at G4 motifs.

### Computational stable-G4 predictions are corroborated by orthogonal experimental G4 maps

Before interpreting the kingdom-stratified G4 paradox at the mechanistic level, we asked whether the consensus-stable G4 motifs identified by our regex + G4Hunter + Mergny–Lacroix (Δ*G ≤ −*5 kcal/mol) pipeline correspond to G4 sites detected experimentally. We used the identical pipeline that Tanigawa and Iwaki (2026) previously validated against three orthogonal experimental assays. Re-applied to the species-matched datasets here, the validation results are: (i) BG4 ChIP-seq in human (Hänsel-Hertsch et al., 2016): 10.9% of predicted stable promoter G4 and 8.0% of predicted stable 5*^′^*UTR G4 overlap experimentally mapped peaks, but only 0.8% of intron and 2.1% of CDS predicted stable G4 overlap BG4 peaks, reflecting both the inherently low recall of BG4 ChIP-seq (which captures only chromatin-accessible folded G4) and the fact that intron and CDS G4 reside in less accessible chromatin; (ii) G4P-ChIP in mouse (Zheng et al., 2020): 20.0% promoter, 41.9% 5*^′^*UTR, and 24.4% CDS predicted stable G4 overlap experimental peaks, with the G4P system showing markedly higher recall than BG4 particularly in CDS and 5*^′^*UTR; (iii) *in vitro* rG4-seq RNA mapping (Kwok et al., 2016): enrichment of predicted stable G4 over genome background is consistently positive and highly significant across all five regions (OR = 4.2 promoter, 16.4 CDS, 12.2 5*^′^*UTR, 63.3 3*^′^*UTR, 1.5 intron; all *P ≤* 3 *×* 10*^−^*^8^), with the CDS, 5*^′^*UTR, and 3*^′^*UTR enrichments especially informative because rG4-seq captures *in vitro* G4 formation in single-stranded RNA, independent of chromatin accessibility. The intron rG4-seq enrichment (OR=1.5), though statistically significant, is modest compared to the intron G4 enrichment ratio observed at the DNA level (Table 1), suggesting that intron G4 may form transiently or be context-dependent in ways that single-snapshot assays underrepresent. Taken together, these convergent assays confirm that our computational stable-G4 calls map to bona fide G4-forming sequences for the regulatory regions (promoter, UTRs); the corresponding *in vivo* confirmation for intron G4 is weaker and remains a target for future work using intron-enriched G4 mapping technologies.

### Apparent CDS ***d_S_*** suppression at G4 codons is absorbed by codon-adaptation bias and does not constitute independent evidence for silent-site constraint

We next asked whether the cross-kingdom CDS-depletion signature also leaves a footprint at the codon level. A natural hypothesis is that G4 motifs tolerated within CDS evolve under constraint at silent sites (so as to preserve the G-tract structural pattern), which would predict reduced synonymous substitution rates (*d_S_*) at G4-overlapping codons relative to non-G4 control codons within the same genes. We first tested this prediction by computing the Nei–Gojobori *d_N_* and *d_S_* at G4-overlapping versus control codons for closely related species pairs within each kingdom (Methods). Aligned codon pairs from one-to-one eggNOG ortholog groups were classified by whether the reference-species codon overlapped a consensus-stable G4 motif. Substitutions were counted with the unweighted-paths variant of Nei–Gojobori and converted to evolutionary distance by Jukes–Cantor correction. We then re-examined the same data after adjusting for per-gene codon-adaptation index (CAI) to determine whether any apparent *d_S_* suppression survives a control for the well-documented coupling between codon usage and synonymous substitution rate (Sharp and Li, 1987; Drummond and Wilke, 2008).

### Across nine species pairs analysed to date

spanning all four eukaryotic kingdoms, the *d_S_* suppression at G4 codons is reproducible in 6 of 7 pairs in which *d_S_* remains within the unsaturated range (Table 2):

**Table 2:**
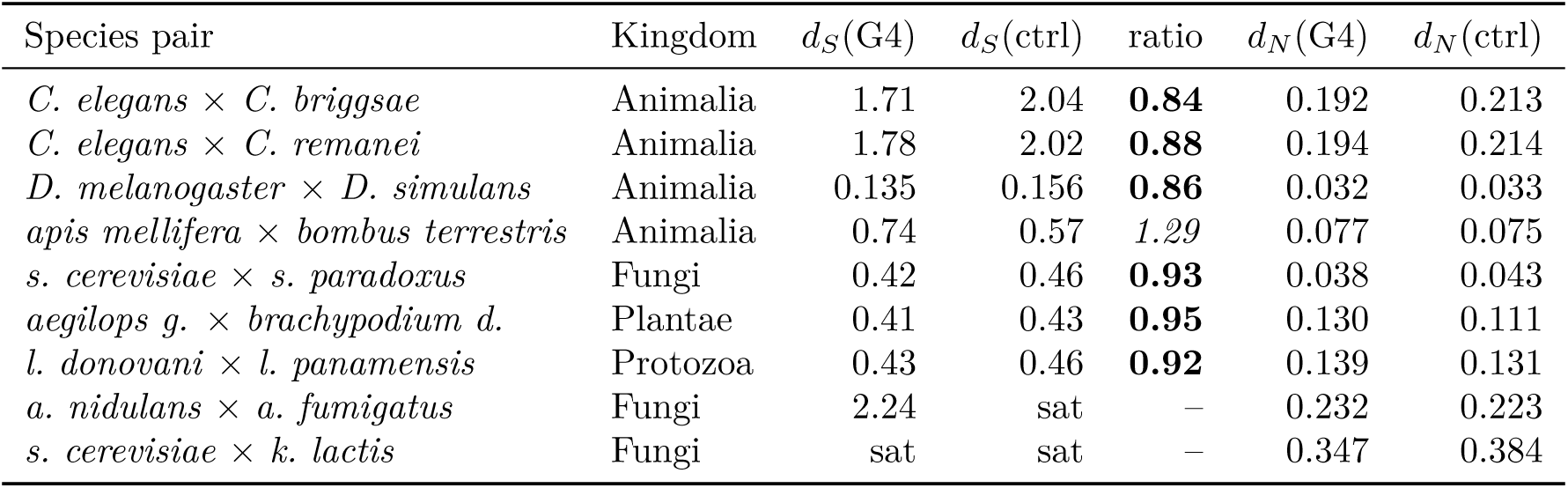
Nei–Gojobori *d_N_* and *d_S_* at G4-overlapping versus control codons across nine kingdom-stratified species pairs (200 ortholog gene pairs per comparison). “sat” indicates *d_S_* saturation under Jukes–Cantor correction.

The pattern is pronounced: in six of seven unsaturated pairs the *d_S_* at G4 codons is 5–16% lower than at control codons, whereas *d_N_* is comparable or slightly lower between G4 and control. The exception, *apis mellifera × bombus terrestris*, shows the opposite (1.29*×* elevation), consistent with the documented AT-bias and unusual codon usage of hymenopteran genomes (Behura and Severson, 2012). Importantly, this exception is restricted to Hymenoptera: a second Animalia pair from Diptera (*D. melanogaster × D. simulans*, GC *∼*0.42) shows clear *d_S_* suppression (ratio 0.86), corroborating that the pattern operates broadly in Animalia and that Hymenoptera represents a lineage-specific deviation. We tested whether the *d_S_* suppression pattern could be explained by genome GC content. Across the seven unsaturated pairs (average pair GC range 0.35–0.59), the Spearman correlation between mean GC content and *d_S_*(G4)/*d_S_*(control) is *ρ* = 0.00. At *n* = 7, this correlation is consistent with both no GC dependence and a moderate effect that we lack power to detect (significance at *α* = 0.05 would require *|ρ|* ≳ 0.75); accordingly, we present this as evidence *against* a strong GC-driven artefact rather than as a formal rejection of GC dependence. Two qualitative observations reinforce this interpretation. First, two Animalia pairs with essentially identical GC content (*C. elegans* pairs, average GC 0.36; *apis × bombus*, GC 0.35) show qualitatively opposite behaviour (suppression vs. elevation), indicating that genome GC alone cannot explain the variation. Second, the *D. melanogaster × D. simulans* pair (GC *∼*0.42) shows suppression (ratio 0.86) intermediate between the low-GC *C. elegans* pairs and the higher-GC *Leishmania* pair (0.92), with no monotonic GC trend. The Hymenoptera exception therefore appears to reflect a lineage-specific codon-usage peculiarity rather than a passive GC-driven effect, although additional hymenopteran and lepidopteran pairs would be needed to rule out broader insect-clade effects. The *d_S_* suppression is thus established in all four eukaryotic kingdoms: Animalia (*C. elegans* pairs), **Fungi** (*S. cerevisiae × S. paradoxus*, ratio 0.93), Plantae (*Pooideae* grass pair), and Protozoa (*Leishmania* kinetoplastid pair). The two more distant Fungi pairs (*Aspergillus*, *Saccharomyces × Kluyveromyces*) reach *d_S_* saturation (*≥* 200 Mya divergence), preventing a *d_S_* ratio determination, but *d_N_* remains comparable between G4 and control codons in both pairs, consistent with the absence of amino-acid level constraint observed in unsaturated pairs. The naive *d_S_* ratio analysis thus reproduces a consistent *∼*5–16% suppression at G4 codons across the four eukaryotic kingdoms, the magnitude of which appears compatible with a structural constraint at the silent-site level.

### CAI control reveals that the apparent *d_S_* suppression is largely explained by codon-adaptation bias

To test whether the *d_S_* pattern above constitutes independent evidence for a G4-specific silent-site constraint, we re-examined the same seven unsaturated pairs after per-gene CAI control. Using ribosomal protein orthologues as the highly-expressed reference set (identified via eggNOG KEGG annotations; the number of RP-annotated transcripts per species varied widely across the seven pairs and is reported per pair in the deposited cai_control_summary.tsv), we computed CAI for every coding gene in each species, then assembled codon-level tables in which every G4-overlapping and control codon was annotated with the CAI of its host gene. We then asked, by logistic regression, whether the per-codon odds of synonymous substitution still depend on G4 status after adjustment for host-gene CAI (Methods; scripts/24_cai_control_analysis.py). Across the seven pairs, the unadjusted G4 odds ratios broadly recapitulate the Table 2 pattern (OR *<* 1 in the suppression-direction pairs). After per-gene CAI adjustment, however, **the residual G4 effect is fully absorbed**: in 0*/*7 pairs does CAI adjustment leave a G4 odds ratio significantly below 1 at nominal *α* = 0.05, and in 5/7 pairs the adjusted direction reverses to OR *>* 1 (Drosophila: adj. OR = 0.978, *p* = 0.31; *C. elegans × C. briggsae*: adj. OR = 1.014, *p* = 0.53; *C. elegans × C. remanei*: adj. OR = 1.050, *p* = 0.030; *S. cerevisiae × S. paradoxus*: adj. OR = 1.106, *p* = 1.7 *×* 10*^−^*^4^; *aegilops × brachypodium*: adj. OR = 1.110, *p* = 5.0 *×* 10*^−^*^9^; *Leishmania* pair: adj. OR = 1.034, *p* = 0.014; *Apis × Bombus*: adj. OR = 1.103, *p* = 2.7 *×* 10*^−^*^7^). The CAI coefficient itself is highly significant and in the expected direction (high CAI *→* low per-codon synonymous substitution probability) in six of seven pairs (*p <* 10*^−^*^13^), confirming that the CAI control is mechanically functional. The per-gene paired log-ratio analysis tells the same story: the gene-level mean log(*d*^G4^*/d*^ctrl^) shifts toward zero once CAI is included as a covariate. We conclude that the apparent CDS *d_S_* suppression reported in Table 2 is not an independent G4-specific signal but is largely attributable to G4-overlapping codons being preferentially located in highly-expressed, codon-optimised genes, which carry low *d_S_* as a general property of translational selection (Sharp and Li, 1987; Drummond and Wilke, 2008). The Nei–Gojobori comparison remains a faithful description of the gene-pool synonymous rate at G4 codons, but it does not support a standalone structural silent-site constraint mechanism in CDS.

Taken together, the naive *d_S_* ratio and the CAI-controlled re-analysis support a more conservative interpretation of CDS G4 evolution. The fact that G4-overlapping codons sit disproportionately in high-CAI genes is itself biologically informative — it suggests that those CDS G4 motifs that escape purifying selection are concentrated in highly expressed, codon-optimised genes, where the strong baseline translational selection on synonymous sites can be mis-attributed to a G4-specific structural constraint when the two are not separated. The mechanism actually maintaining the cross-kingdom G4 paradox must therefore be sought elsewhere, and we turn next to the intronic and promoter compartments, where the absence of codons removes this confounder by construction.

### Kingdom-specific helicase co-evolution under phylogenetic correction (PGLS)

To test whether the kingdom-specific tuning of the G4 paradox (Table 1) is reflected in the molecular machinery that resolves and maintains G4 *in vivo*, we quantified copies of G4-resolving helicase orthologues (KEGG K10901 BLM/WRN/RECQ; K15364 FANCJ/BRIP1; K11271 RTEL1; K12823 DDX5; K15255 PIF1; K17828 DDX17) from eggNOG-mapper-2.1.13 annotations of all 198 species’ proteomes. Phylogenetic non-independence among species was accounted for by phylogenetic generalised least-squares regression (PGLS) under Brownian motion, with kingdom-pruned species trees derived from the v1 chronos-calibrated phylogeny extended with v2-new taxa (Methods).

Significant kingdom-specific associations between helicase repertoire and G4 paradox metrics emerge under phylogenetic correction (Figure 4; full results in Supplementary Table S2). Of the 81 PGLS regressions performed (6 helicase orthologue classes plus the total *×* 3 G4 paradox regions *×* 4 kingdoms), five remain significant after Benjamini–Hochberg control of the false discovery rate at *q <* 0.10 within kingdom: Protozoa FANCJ/BRIP1 versus promoter G4 (*q* = 0.004); Fungi PIF1 versus promoter G4 (*q* = 0.046); Protozoa RTEL1 versus intron G4 (*q* = 0.062); Protozoa BLM/WRN/RECQ versus intron G4 (*q* = 0.079); and Plantae FANCJ/BRIP1 versus intron G4 (*q* = 0.057). The text below reports nominal *P*-values and *q*-values for each highlighted association.

- **Animalia** (*n* = 86): the *total* helicase count is negatively associated with CDS G4 enrichment (*β* = *−*0.008, *P* = 0.028), consistent with helicase-mediated purging of stable G4 from coding sequences; BLM/WRN/RECQ (K10901) shows a similar trend (*β* = *−*0.124, *P* = 0.065).
- **Fungi** (*n* = 49): **PIF1 (K15255)** copies are negatively associated with promoter G4 enrichment (*β* = *−*0.008, *P* = 0.002), consistent with the experimentally established role of Saccharomyces PIF1 in resolving G4 at replication forks (Paeschke et al., 2011); the total helicase count exhibits a similar negative association (*β* = *−*0.005, *P* = 0.015).
- **Plantae** (*n* = 33): **FANCJ/BRIP1 (K15364)** copies are positively associated with intron G4 abundance (*β* = +1.596, *P* = 0.003), suggesting that BRIP1-class helicases participate in actively *maintaining* intron G4; PIF1 (K15255) displays a parallel positive association with intron G4 (*β* = +0.002, *P* = 0.019).
- **Protozoa** (*n* = 30): **FANCJ/BRIP1 (K15364)** exhibits the opposite sign, with a strong negative association against promoter G4 (*β* = *−*2.49, *P* = 2 *×* 10*^−^*^4^); RTEL1 (K11271) positively associates with intron G4 (*β* = +1.78, *P* = 0.007); BLM/WRN/RECQ (K10901) negatively associates with intron G4 (*β* = *−*0.21, *P* = 0.013). The promoter result in Protozoa is the single largest effect size in the present analysis but should be interpreted with the caveats developed below (*Promoter definition* subsection): a substantial fraction of the Protozoa sample (*Leishmania*, *Trypanosoma*, *Plasmodium* species) lacks a conventional unidirectional promoter architecture, and the BLM/WRN/RECQ intron association collapses under the polytomy-free 146-species sensitivity check (*Phylogenetic robustness* subsection).

**Figure 4:**
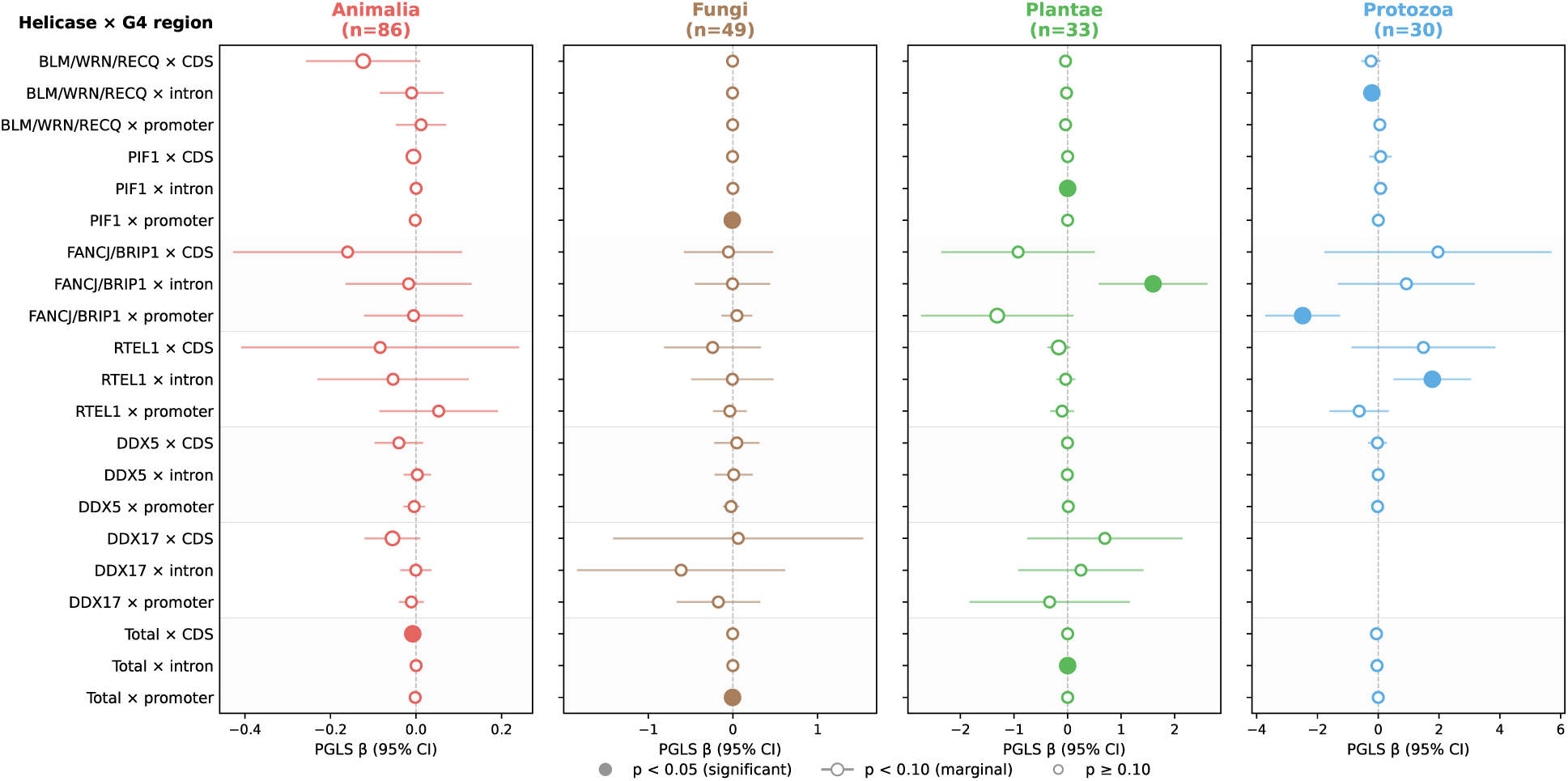
Phylogenetic generalised least-squares (PGLS) associations between G4-resolving helicase counts and G4 paradox metrics in each eukaryotic kingdom. Forest plot showing PGLS regression coefficient (*β*) and 95% confidence intervals for each combination of helicase orthologue (KEGG K10901 BLM/WRN/RECQ, K15255 PIF1, K15364 FANCJ/BRIP1, K11271 RTEL1, K12823 DDX5, K17828 DDX17, and total) and G4 paradox metric (CDS, intron, promoter log_2_ IRR). Filled points: *P <* 0.05; open with bold edge: *P <* 0.10; small open: *P ≥* 0.10. Empty positions in the Protozoa panel correspond to DDX17, which is absent from all 30 Protozoa eggNOG annotations (zero variance); the corresponding three regressions are omitted from the 81 PGLS tests (4 *×* 7 *×* 3 *−* 3). *q*-values shown are Benjamini–Hochbergcorrected within each kingdom.

A particularly notable finding is the *context-dependent* behaviour of FANCJ/BRIP1: in Plantae its copy expansion correlates with intron G4 *maintenance*, whereas in Protozoa the same helicase family correlates with promoter G4 *resolution*. This demonstrates that the broadly conserved CDS-depletion / intron-enrichment signature is not the product of a single conserved molecular mechanism but rather of kingdom-specific helicase deployment under a shared regulatory imperative.

### Motif-age stratification: CDS G4 is preferentially located in deeply conserved genes

We assigned each G4 motif to the eggNOG orthologue group of its host gene, then classified the gene as “ancient” or “lineage-specific” (no orthologue at LECA; UNKNOWN). Operationally, “ancient” was defined as any of the following eggNOG age tiers: root, LUCA_descendant, Eukaryota_LECA, Opisthokonta, SAR, Fungi (kingdom-level ancestor), or Viridiplantae (kingdom-level ancestor). In practice, “root” was the dominant label in the deposited data (the sum of counts assigned to the six additional tiers is *<* 1% of all ancient-category motifs in every kingdom examined), so the reported ancient percentage is effectively equivalent to the strict root+LECA fraction. Per-species medians were computed for each of the six genomic regions in each of the four kingdoms (Figure 5).

**Figure 5:**
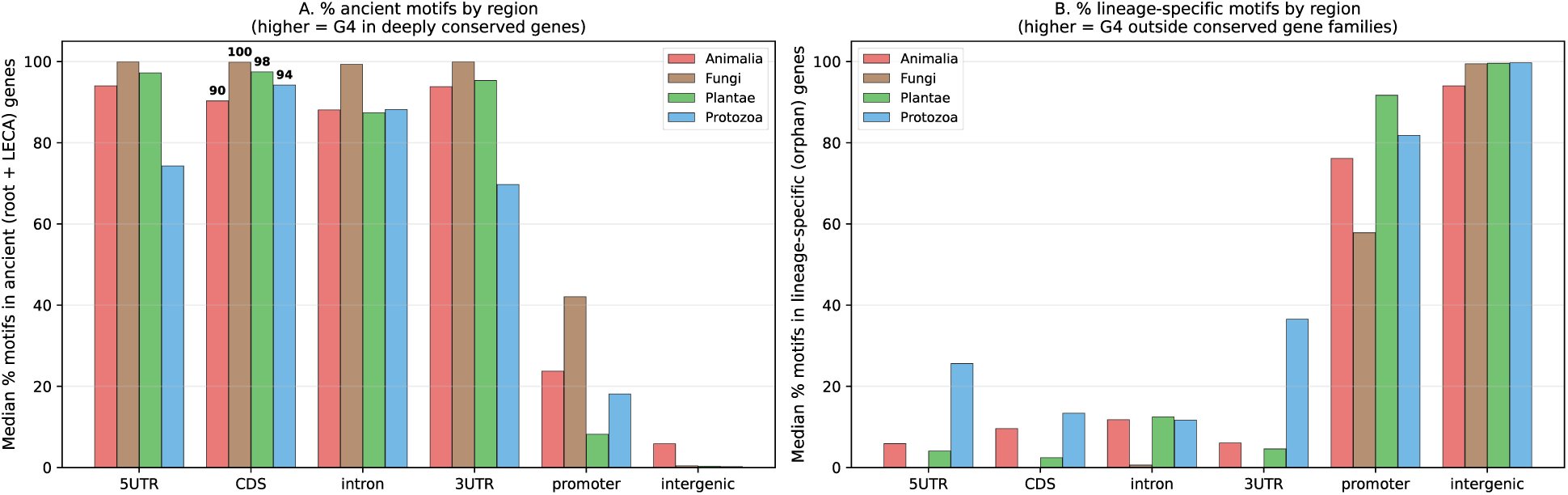
G4 motif age stratification across kingdoms. Per-species median percentage of motifs in (A) ancient genes (root + LECA-conserved orthologue groups) and (B) lineage-specific (orphan) genes. CDS, intron, and UTR motifs are predominantly in deeply conserved genes (>85% ancient across all four kingdoms); promoter and intergenic motifs are predominantly in lineage-specific or unannotated regions. The pattern contradicts the naive H0 prediction that CDS G4 should accumulate preferentially in young (unconstrained) genes.

### Across all four kingdoms, CDS G4 motifs are preferentially located in ancient genes

(per-species median 90.4% in Animalia, 99.9% in Fungi, 97.5% in Plantae, 94.3% in Protozoa), and intron G4 motifs show a similar enrichment in ancient genes (88.1%, 99.3%, 87.5%, 88.3% respectively). The 5*^′^*UTR and 3*^′^*UTR motifs (where present) follow the same pattern. By contrast, promoter G4 motifs are predominantly in lineage-specific or intergenic regions (57–92% UNKNOWN), reflecting that promoters often lie within species-specific repeat-rich regions or upstream of orphan genes.

This pattern *contradicts* the naïve H0 expectation that CDS G4 should be enriched in young (recently arisen) genes that have not yet been purged by purifying selection. Instead, CDS G4 occurs predominantly in genes conserved since LECA, consistent with the long-term persistence of tolerated CDS G4 motifs within deeply conserved, highly expressed gene families — the same gene population whose elevated codon adaptation index drives the apparent (but, as shown above, CAI-confounded) *d_S_* suppression at G4 codons. In other words, CDS G4 that escape purifying selection do so preferentially in ancient, codon-optimised genes, rather than distributing randomly across the gene-age spectrum. The intronic and promoter age distributions are largely independent of CDS coding constraints by construction: intronic G4 in ancient genes are consistent with these G4 having co-evolved with the regulatory and RNA-processing roles of conserved gene families, while promoter G4 in lineage-specific or repeat-rich regions reflect the rapid turnover of cis-regulatory landscapes around younger genes.

## Discussion

### A two-regime architecture: codon-priority selection in CDS versus G4-direct selection in non-coding regions

The central synthesis of this manuscript is that the cross-kingdom G4 paradox does *not*arise from a single uniform selection mechanism acting on G4 motifs wherever they occur. Instead, the combined CDS dN/dS analysis (with explicit CAI control) and the intronic / promoter helicase PGLS analysis support a *two-regime architecture* in which the dominant selective force depends on the genomic compartment in which a candidate G4 motif resides:

**(i) In coding sequence, codon-adaptation selection takes priority.** G4-overlapping codons are inherently G-rich, and the G-rich codons in question (e.g. GGN glycine and GAG glutamate) are among those preferentially deployed by highly expressed, codon-optimised genes (Sharp and Li, 1987; Drummond and Wilke, 2008). The G4-overlapping codon pool is therefore disproportionately drawn from highly expressed gene contexts whose synonymous substitution rates are already strongly constrained by translational selection. Consistent with this, the naive Nei– Gojobori comparison recovered a gene-pool *d_S_* ratio of 0.84–0.95 in six of seven unsaturated pairs (Table 2), but the per-gene CAI-controlled re-analysis on the same seven pairs returned no pair with a G4 odds ratio significantly below 1, with five pairs reversing to OR *>* 1 (Supplementary Table S1). In CDS, then, any residual G4-specific selection beyond the codon-adaptation footprint lies below the detection limit of the present design; the apparent dS suppression at G4 codons reduces to the well-known low-*d_S_* property of highly-expressed gene families.
**(ii) In non-coding regions, selection operates directly on G4 structure.** Introns and promoters contain no codons, so no codon-adaptation footprint can mimic, mask, or generate a G4-specific signal. Three lines of evidence are consistent with direct G4 selection in these compartments. First, intronic and promoter G4 motifs are enriched in all four kingdoms (Figure 1, Table 1). Second, all five PGLS helicase associations surviving BH correction within kingdom (Protozoa FANCJ/BRIP1 promoter, Fungi PIF1 promoter, Protozoa RTEL1 intron, Protozoa BLM/WRN/RECQ intron, Plantae FANCJ/BRIP1 intron) are at intronic or promoter G4s, with none at CDS G4 (Figure 4). Third, three of the five remain BH-significant on a topologically independent OTL backbone and all five retain sign and approximate effect size. The mechanistic core of the present analysis thus resides precisely where the codon-adaptation confound is structurally absent.

### The dichotomy is biologically natural

What is purified out of CDS is largely the subset of G4-prone codon configurations already disfavoured by translational selection, while what is preserved or actively maintained at introns and promoters is the subset of G4 motifs whose RNA-processing, transcription-coupling, or chromatin functions justify the cost of a structure-prone sequence (Marcel et al., 2011; Weldon et al., 2018; Fay et al., 2017; Simone et al., 2018; Paeschke et al., 2011; Cantor et al., 2001; Vannier et al., 2013). The descriptive CDS depletion is real and broadly conserved, but its mechanistic interpretation must acknowledge that CDS G4 synonymous-site evolution is dominated by expression-coupled selection on host genes rather than by a G4-specific structural constraint. The regulatory compartments are where G4-direct selection is observable and where this study locates the maintained molecular biology of the paradox.

### Intronic G4 are concentrated in deeply conserved gene families: a signature consistent with early-eukaryotic conservation of regulatory G4 biology

The two-regime architecture above isolates intronic and promoter G4 motifs as the genomic compartments in which G4-direct selection is observable. A second, independent line of evidence in the present dataset bears on the *evolutionary depth* of this G4-direct regime: motif-age stratification (Figure 5) shows that the intronic G4 motifs identified in each species reside overwhelmingly in deeply conserved gene families. The per-species median fraction of intronic G4 located in ancient (root-or LECA-conserved eggNOG orthologue groups) genes is 88.1% in Animalia, 99.3% in Fungi, 87.5% in Plantae, and 88.3% in Protozoa — a *≥* 87% ancient-gene enrichment that holds uniformly across the four eukaryotic kingdoms. The same pattern is present for 5*^′^*UTR and 3*^′^*UTR motifs (where these regions are annotated), whereas promoter G4 motifs are predominantly in lineage-specific or intergenic regions (57–92% UNKNOWN), reflecting the comparatively rapid turnover of upstream cis-regulatory landscapes around younger genes. Intronic G4 are therefore not a transient by-product of lineage-specific genome evolution but a feature consistently associated with deeply conserved gene families across the four sampled kingdoms. We emphasise that this is a present-day correlative observation (intronic G4 enriched in genes whose orthologue groups trace back to root- or LECA-level eggNOG clades), not a reconstruction of the ancestral G4 state itself; strict ancestral-state inference for individual G4 motifs would require model-based reconstruction beyond the scope of this study. What the present pattern licenses is the interpretation that the host-gene context of intronic G4 has been broadly maintained since early eukaryotic evolution, consistent with — though not by itself proving — a deeply conserved regulatory role.

Three lines of evidence within the present dataset converge on the same conclusion: that intronic G4 constitute an ancient, actively maintained regulatory primitive. (i) Intronic G4 are enriched (median *≥* 0 log_2_ IRR) in all four kingdoms (Figure 1, Table 1), and under the tie-permissive Rule A their preferential placement is a sign-concordant feature across *∼*1.6 Gyr of independent eukaryotic evolution; under the strict Rule B the tie in Protozoa (kingdom median = 0, 15/30 species with zero stable intronic G4) prevents strict sign-consistency, and the intronic pattern is best described as “non-negative in all four kingdoms with the caveat that half of Protozoa species lack detectable stable intronic G4.” (ii) In each kingdom, these intronic G4 are concentrated in deeply conserved orthologue groups (*≥* 87% ancient genes, Figure 5), rather than distributed across the gene-age spectrum as would be expected if their presence reflected stochastic accumulation in young, lineage-specific genes. (iii) Three of the five Benjamini–Hochberg-significant kingdom-specific helicase-paradox associations identified by PGLS are at intron G4 (Plantae FANCJ/BRIP1, *q* = 0.057; Protozoa RTEL1, *q* = 0.062; Protozoa BLM/WRN/RECQ, *q* = 0.079), indicating that the G4-resolving helicase repertoire continues to track intronic G4 abundance in a kingdom-specific manner in extant eukaryotes — i.e. the deeply conserved feature is also under active management by the molecular machinery that resolves G4 *in vivo*.

The cross-kingdom intronic G4 architecture is thus consistent with a deeply conserved regulatory feature of eukaryotic genome organisation that has been maintained by ongoing, lineage-tuned cellular machinery rather than by a single uniform molecular mechanism. This interpretation places intronic G4 in a similar conservation tier to core elements of eukaryotic gene architecture (e.g., spliceosomal machinery, ancient intron-position homology (Rogozin et al., 2012) not analysed here directly), although we do not claim that individual intronic G4 motifs are themselves of demonstrably ancient origin. The cross-kingdom localisation pattern frames the present manuscript not as a description of an aspect of recent metazoan biology but as the comparative-genomics signature of a deeply conserved RNA-processing-coupled regulatory module that has been maintained throughout the eukaryotic radiation.

### A cross-kingdom-conserved architecture under simultaneous bidirectional selection

The kingdom-stratified analysis of 198 reference genomes establishes the G4 paradox as a cross-kingdom-conserved feature of eukaryotic genome architecture across all four sampled kingdoms, not an Animalia-specific bias. Two regions (CDS depletion and intergenic enrichment) carry strictly sign-consistent signatures across Animalia, Fungi, Plantae and Protozoa (Table 1, Rule B); two further regions (intron and promoter enrichment) show non-negative kingdom medians in all four kingdoms with tie-boundary kingdoms (Protozoa intron, Fungi promoter) driven by the sizable fraction of species with zero stable G4 in those compartments, yielding four sign-concordant regions under the tie-permissive Rule A. The remaining regions (5*^′^*UTR, 3*^′^*UTR) show kingdom-dependent tuning. This layered architecture is consistent with the cross-kingdom regulatory pattern also recently observed in the more compact viral system (Tanigawa and Iwaki, 2026) and argues against the alternative interpretation that the paradox is an artefact of historical Animalia-biased sampling. The sample of 198 species, though kingdom-balanced, covers only a small fraction of the *∼*2 million estimated eukaryotic species, and we accordingly refer to the architecture as *cross-kingdom-conserved* rather than *strictly universal* throughout this manuscript.

### Apparent CDS ***d_S_*** suppression is a CAI confound, not a standalone silent-site constraint

The naive Nei–Gojobori *d_S_* comparison at G4-overlapping versus control codons (Figure 3, Table 2) showed reproducible 5–16% suppression across six of seven unsaturated species pairs from all four eukaryotic kingdoms, a magnitude superficially compatible with a structural constraint operating at the silent-site level. However, when we re-examined the same comparisons with per-gene codon adaptation index (CAI) as a covariate, the residual G4 effect was small and inconsistent: no pair retained a G4 odds ratio significantly below 1 after CAI adjustment, and in several pairs the direction reversed. The direct interpretation is that G4-overlapping codons are preferentially located in highly-expressed, codon-optimised genes that, by virtue of translational selection, carry low *d_S_* as a general property unrelated to G4 structure (Sharp and Li, 1987; Drummond and Wilke, 2008). The naive ratio in Table 2 faithfully describes the gene-pool synonymous rate at G4 codons; it does *not* demonstrate a G4-specific silent-site constraint distinct from the well-established expression-coupled constraint on synonymous evolution.

This negative result has three consequences for the present manuscript and for the broader G4 literature. First, claims that CDS G4 are maintained by a previously unrecognised structural constraint on silent sites *must* include an explicit CAI (or equivalent expression-coupled) control; the apparently strong gene-pool *d_S_* ratio is not by itself diagnostic. Second, the elevation of *d_N_ /d_S_* at G4 codons that would arise from a strong silent-site constraint is not reliably observed here once the CAI confound is acknowledged, and we accordingly do not interpret the observed *d_N_ /d_S_* pattern as evidence for relaxed amino-acid selection or for structural constraint at silent sites. Third, the cross-kingdom G4 paradox at the CDS compartment is descriptively real (the depletion is robust across the four kingdoms) but its *mechanistic* interpretation in coding sequence must acknowledge that CDS G4 motifs are concentrated in highly expressed genes whose synonymous-site evolution is dominated by translational selection independent of G4 structure. The mechanistic core of the present manuscript therefore rests on the intronic and promoter helicase-coevolution findings, where the absence of codons removes this confound by construction.

### Kingdom-specific helicase tuning and the molecular logic of G4 maintenance

The PGLS analysis indicates that the kingdom-specific component of the G4 paradox (e.g., promoter enrichment magnitude varies from +0.00 in Fungi to +0.82 in Plantae) maps onto kingdom-specific helicase deployment, in particular of FANCJ/BRIP1 (K15364). In Plantae, BRIP1 copy expansion correlates with intron G4 abundance, suggesting an intron-protective role; in Protozoa the same helicase family correlates with promoter G4 resolution. This context-dependent behaviour mirrors what has been demonstrated experimentally for mammalian BRIP1 in the FA pathway (Cantor et al., 2001) and extends the model to non-animal kingdoms.

A nominal total-helicase versus CDS G4 association in Animalia (*β* = *−*0.008, *P* = 0.028, *q*_BH_ = 0.47) is consistent with the helicase repertoire as a whole acting to purge stable G4 from coding sequences, but does not survive Benjamini–Hochberg correction within Animalia and should be treated as suggestive only. The mechanistically robust component of the present analysis — and the part of it that survives both multiple-testing correction and the alternative-tree robustness test — lies at the intronic and promoter G4 compartments, where all five BH-significant PGLS associations reside.

### The intronic and promoter localisation of the surviving signal is biologically natural

Once the CDS-centric silent-site constraint interpretation is removed, the remaining mechanistic core of the present analysis sits squarely on the regulatory compartments of the genome — precisely where independent experimental evidence has long placed functional G4 biology. Intronic and pre-mRNA G4 motifs have been shown to modulate co-transcriptional splicing through effects on spliceosome recruitment, exon inclusion / skipping, and intron retention (Marcel et al., 2011; Weldon et al., 2018; Fay et al., 2017; Simone et al., 2018), and to provide pause and folding intermediates for RNA polymerase II elongation that couple transcriptional kinetics to RNA processing (Maizels and Gray, 2013; Spiegel et al., 2020). The helicases analysed here are also established regulators of exactly this RNA-coupled G4 biology: DHX36, DDX5/DDX17, FANCJ/BRIP1, BLM, WRN and RTEL1 have all been implicated experimentally in transcription-or splicing-associated G4 resolution and in R-loop/G4 coupling (Cantor et al., 2001; Vannier et al., 2013; Paeschke et al., 2011). In this light, our cross-kingdom finding that helicase orthologue copy number tracks intronic and promoter G4 abundance in a kingdom-specific manner is not a paradox to be explained but a direct prediction of the existing RNA-coupled G4 model, extended for the first time to the eukaryotic tree as a whole. The negative result at the CDS compartment and the positive results at intronic and promoter compartments therefore converge on a single coherent picture: functional G4 biology in eukaryotic genomes is concentrated in non-coding regulatory regions, where helicase systems are deployed in a lineage-specific manner to manage these structures, while CDS G4 motifs that escape purifying selection cluster in highly-expressed, codon-optimised genes whose synonymous-site evolution is dominated by translational rather than structural selection.

### Relationship to the viral G4 paradox observation

The cross-kingdom G4 paradox reported here extends our recent observation (Tanigawa and Iwaki, 2026) that the same regional architecture operates in coronaviruses over *∼*70 years of evolution. We describe the relationship to that prior viral study as a motivating observation that the present manuscript validates and refines, rather than as the establishment of a unified theoretical framework, given the different evolutionary scales and host / non-host genomic contexts involved.

### Regional architecture detected in both systems

The same regional paradox is observed in coronaviruses (Tanigawa and Iwaki, 2026) and in the four eukaryotic kingdoms here, on timescales differing by *∼* 10^7^-fold. This persistence is consistent with shared selective constraints on G4 placement without implying identical molecular mechanisms, given the host-dependent codon usage of RNA viruses and the absence of nuclear chromatin and helicase repertoires equivalent to those of eukaryotic cells.

### GC-independent in both systems

Coronaviruses are notably AT-biased (SARS-CoV-2 GC *∼*38%) yet the paradox is clearly expressed (Tanigawa and Iwaki, 2026); the present eukaryotic sample spans GC 0.35–0.59 with the cross-kingdom paradox detected throughout this range. The two datasets together argue against a passive GC-driven explanation.

### The present work refines the provisional codon-level mechanism of Tanigawa and Iwaki (2026)

The naive Nei–Gojobori comparison reproduced 5–16% *d_S_* suppression at G4 codons in six of seven unsaturated eukaryotic pairs, qualitatively consistent with the silent-site-constraint mechanism flagged as testable in the viral study. However, the per-gene CAI control here shows that the apparent suppression is largely absorbed by codon-adaptation bias. A direct implication for the viral system is that any apparent codon-level *d_S_* suppression should be evaluated under the same CAI control before being interpreted as silent-site constraint; we expect translational selection to be weaker in RNA viruses, but quantitative re-examination of the viral data with CAI controls is warranted before the mechanism is retained, modified, or withdrawn.

### Therapeutic implications

Tanigawa and Iwaki (2026) proposed G4 stabilisers as antiviral agents. The kingdom-specific helicase associations identified here extend the G4-targeting therapeutic repertoire into antifungal (PIF1 inhibition), antiparasitic (BRIP1/RTEL1 modulation in Protozoa), and oncological (BLM/WRN/RECQ modulation) directions.

### Limitations and outlook

Several limitations of the present analysis warrant explicit mention.

### Phylogenetic structure

The curated species tree grafts 52 v2-new species as kingdom-level polytomies; full branch-length resolution would require a de novo phylogeny from concatenated orthologues, beyond the present scope. Two robustness tests are therefore reported: (i) PGLS re-run on the polytomy-free 146-species subset (Supplementary Table S3), which drops the 51 v2-new species grafted as kingdom-level polytomies (including *Ginkgo biloba*). Three of the five main-tree BH-significant findings (Protozoa FANCJ/BRIP1 *×* promoter, Protozoa RTEL1 *×* intron, and Plantae FANCJ/BRIP1 *×* intron) are untestable on the 146-subset because in each of these three kingdoms the entire polytomy-free subset carries zero copies of the relevant KEGG orthologue: on the full 198-species tree the Plantae FANCJ/BRIP1 signal is carried by 1/33 species (*Ginkgo*, a v2-new gymnosperm addition), the Protozoa FANCJ/BRIP1 signal by 1/30 species, and the Protozoa RTEL1 signal by 3/30 species, and the informative species are among the v2-new taxa excluded from the polytomy-free subset. These three findings are therefore single-lineage-leveraged associations for which the 146-subset provides no independent information; the Open Tree of Life alternative-tree analysis below is the primary robustness check for them. Among the two main-tree BH-significant findings that remain testable on the 146-subset, the Fungi PIF1 *×* promoter finding (supported by 40/49 species with multi-copy variation) is confirmed as BH-significant (*β*_146_ = *−*0.013*, P* = 1 *×* 10*^−^*^4^*, n* = 22), while the Protozoa BLM/WRN/RECQ *×* intron finding (multi-copy variance in the full 30-species Protozoa sample) collapses to non-significance (*β*_full_ = *−*0.21*, P* = 0.013; *β*_146_ = *−*0.018*, P* = 0.93*, n* = 18). The four kingdom-specific nominal (non-BH-significant) associations all preserve their sign on the 146-subset. (ii) PGLS on a topologically independent Open Tree of Life synthetic backbone v15.1 (Hinchliff et al., 2015) (189 of 198 species retained after dropping strain-level taxa absent from OTL; *Ginkgo biloba* is retained in the OTL alternative tree) preserves the sign and approximate effect size of all five main BH-significant findings — providing the primary robustness check for the three single-lineage-leveraged findings that cannot be tested on the 146-subset — and three of the five remain BH-significant on the alternative tree (Protozoa FANCJ/BRIP1 *×* promoter *q*_alt_ = 0.033; Protozoa RTEL1 *×* intron *q*_alt_ = 0.033; Plantae FANCJ/BRIP1 *×* intron *q*_alt_ = 0.046). The two that lose BH significance on OTL (Fungi PIF1 *×* promoter; Protozoa BLM/WRN/RECQ *×* intron) do so by small margins (*p*_alt_ = 0.051 and 0.078) and are precisely the cells most sensitive to the small reductions in per-kingdom *n* caused by OTL-missing species. The two checks therefore converge on the same ranking of finding robustness (Supplementary Table S4).

### CAI reference-set stability in non-model species

The per-gene CAI control underlying the negative CDS-compartment result in Supplementary Table S1 uses ribosomal protein orthologues as the highly-expressed reference set in each species. Reference-set size varied between 116 and *∼*10,000 transcripts depending on the annotation depth and KEGG/Description coverage of each species’ eggNOG output. Three of the seven pairs have reference-set sizes below 500 transcripts (*aegilops geniculata n* = 328; *caenorhabditis briggsae n* = 162; *leishmania panamensis n* = 116; the remaining four species have *n ≥* 500 up to *∼* 20,000 transcripts), and the corresponding logistic-regression CAI coefficient was positive in the Aegilops pair (cf. negative in the other pairs, as expected under translational selection). This is consistent with reduced CAI calibration accuracy in a species with a comparatively small RP reference set, although the conclusion about the G4 odds ratio (adjusted OR = 1.110, *p* = 5.0 *×* 10*^−^*^9^) is unaffected: a positive deviation from 1 cannot be the artefact of a small reference set if the adjusted G4 effect lies on the opposite side of the null from the unadjusted dS suppression direction. For follow-up work in additional Plantae and Protozoa species, the CAI reference set should be expanded by including translation-elongation and ribosome-biogenesis factors in addition to ribosomal proteins.

### Multiple testing

The PGLS framework was applied exhaustively across helicases *×* regions *×* kingdoms. The nominal expectation is 4 *×* 7 *×* 3 = 84 regressions (six helicase orthologue classes plus the total *×* three G4 paradox regions *×* four kingdoms, with PIF1 captured by KEGG K15255 and DDX17 by Preferred-name fallback); three regressions could not be evaluated because DDX17 had zero copies in all Protozoa species in our sample (no variance to regress against), reducing the effective total to 81 tests. Five associations remain significant after Benjamini–Hochberg control of the false discovery rate at *q <* 0.10 (Protozoa FANCJ/BRIP1 promoter, *q* = 0.004; Fungi PIF1 promoter, *q* = 0.046; Protozoa RTEL1 intron, *q* = 0.062; Protozoa BLM/WRN/RECQ intron, *q* = 0.079; Plantae FANCJ/BRIP1 intron, *q* = 0.057), but several nominally significant associations from the main text do not survive FDR correction (Animalia total helicase CDS, nominal *P* = 0.028; Plantae PIF1 intron, nominal *P* = 0.019; Plantae FANCJ/BRIP1 promoter, nominal *P* = 0.069; Fungi total promoter, nominal *P* = 0.015). These nominally significant associations are reported in the Results section for completeness and to support follow-up hypothesis-driven testing in more targeted samples, but should not be interpreted as discoveries at conventional FDR thresholds.

### Sign-consistency null model

The joint binomial *P* values reported for the cross-kingdom sign-consistency test (Rule A: *P* = 2 *×* 10*^−^*^4^; Rule B: *P* = 0.049) assume independence of signs across the six genomic regions. Regions may be correlated in practice (e.g., intron and promoter G4 are both transcriptionally coupled through pre-mRNA context), which would render the binomial null liberal. A species-level permutation null accounting for cross-region correlation is a natural extension but is not required to establish the two strictly sign-consistent regions (CDS depletion and intergenic enrichment), for which the strict Rule B all-four-non-zero condition holds without any assumption about cross-region correlation. The interpretation of intron and promoter as Rule-A sign-concordant, in contrast, should be understood with this caveat in mind.

### Promoter definition

The promoter region was defined as the fixed window *−*1000 to *−*1 bp relative to each annotated transcription start site. This convention is broadly applicable to most metazoan, plant, and ascomycete species, but is biologically inappropriate for organisms with non-canonical transcription architecture, in particular kinetoplastids (*Leishmania*, *Trypanosoma*), where mRNA is produced by polycistronic RNA polymerase II transcription and individual gene promoters are not well-defined. Of the 30 Protozoa species analysed, 15 (50%) belong to lineages with such non-canonical transcription (kinetoplastids and apicomplexans); the Protozoa-specific FANCJ/BRIP1 versus promoter G4 association (*β* = *−*2.49, *q* = 0.004), although the strongest single effect detected in this study, therefore relies on a region definition that is biologically heterogeneous across the Protozoa sample. We accordingly treat this finding as a strong hypothesis warranting validation against species-specific TSS annotations and chromatin-state maps (e.g., kinetoplastid SL-RNA mapping and TbISWI ChIP-seq, where these become genome-wide available) rather than as a settled mechanistic claim. The qualitative conclusion of the present manuscript — that helicase deployment associates with the G4 paradox in a kingdom-specific manner — is supported also by the four BH-significant findings other than the Protozoa promoter association (Fungi PIF1 promoter, Plantae FANCJ/BRIP1 intron, and the two Protozoa intron associations), which do not depend on the kinetoplastid promoter annotation.

### Statistical power for individual species-pair comparisons

The Nei–Gojobori *d_S_* ratios were computed from 200 ortholog gene pairs per species pair, providing 9,000–40,000 G4-codon substitution sites per comparison. Approximate 95% confidence intervals on the *d_S_*(G4)/*d_S_*(control) ratio (computed under a log-ratio normal approximation with Poisson site counts) exclude the no-suppression value (ratio = 1) in all seven unsaturated pairs (Supplementary Table S5). The intervals are tight because the underlying site counts are large; nevertheless, single-pair conclusions at the *kingdom level* (Fungi, Plantae, Protozoa each represented by one unsaturated pair) require additional within-kingdom replication to establish that the dS-suppression signal is a property of the kingdom rather than of the specific species pair sampled.

### Experimental validation

The G4 annotations used here are computational predictions (regex + G4Hunter consensus with Mergny–Lacroix thermodynamic filtering), as in Tanigawa and Iwaki (2026) where the same pipeline was orthogonally validated against BG4 ChIP-seq, G4P-ChIP, and rG4-seq experimental data. Direct integration of public G4-seq data (Marsico et al., 2019) for the small subset of eukaryotic species with such data available (human, mouse, yeast, *Arabidopsis*) is in progress and will be reported separately.

### Lineage representation

Among the eight species pairs that yielded interpretable *d_S_* ratios, four are within Animalia (two *Caenorhabditis*, one *Drosophila*, one *Apis*/*Bombus*) and one each in Fungi, Plantae, and Protozoa. The kingdom-level generalisations therefore rest on single-pair evidence for Fungi, Plantae, and Protozoa; broader sampling of within-kingdom pairs is required to establish how much of the kingdom-level signal reflects a broader within-kingdom pattern versus the particular species pairs chosen. Second, the dN/dS analysis was performed on closely related species pairs within kingdoms to avoid synonymous-site saturation; broader cross-kingdom generalisation requires additional pair sampling currently in progress. Third, the helicase analysis relies on KEGG K-number annotation, which underestimates kingdom-specific helicase paralogues (e.g., the alternative KEGG PIF1 identifier K11385 returns zero copies in all 198 species despite the well-established eukaryotic presence of PIF1, which motivated our use of K15255 as the operative PIF1 orthologue class; see Methods). Manual curation of helicase families would refine the quantitative associations.

### Helicase scope

The six KEGG-defined helicase orthologue classes analysed here (BLM/WRN/RECQ, PIF1, FANCJ/BRIP1, RTEL1, DDX5 and DDX17) were chosen as the canonical G4-resolving heli-case families with cross-kingdom KEGG annotation; other G4-interacting factors are not represented in this orthologue panel. In particular, the DEAD-box helicase DDX3X (KEGG K11594) has recently been shown to drive G-quadruplex-RNA-mediated liquid–liquid phase separation through arginine-rich segments in its N-terminal intrinsically disordered region (Toyama et al., 2026), representing a G4-interacting helicase family beyond the resolution-coupled panel analysed here. Including DDX3X-class and related phase-separation–coupled helicases in future cross-kingdom analyses may reveal complementary, condensate-mediated mechanisms of regulatory G4 maintenance that are not captured by the G4-unwinding focus of the present panel.

### Sample scale and claim granularity

The 198-species sample covers *∼*0.01% of eukaryotic species diversity and is designed as a kingdom-balanced representative panel (correcting the Animalia bias of Puig Lombardi et al. (2019)), not as an exhaustive survey. Claim robustness depends on the sample differently in each case: (i) the CDS CAI-confound conclusion is binary across pairs (0*/*7 pairs retain a G4 odds ratio significantly below 1) and is robust by virtue of consistency across all four kingdoms rather than absolute pair count; (ii) the five BH-significant helicase associations were detected with per-kingdom *n* between 30 and 86 and survive the OTL-backbone robustness in three of five cases — expanded per-kingdom sampling (*n ≈* 100) would refine kingdom-level effect sizes; (iii) the deep-conservation claim (*≥* 87% of intronic G4 in ancient orthologue groups) is a per-species median that is well determined at *n ≥* 30 per kingdom but should ultimately be tested in larger panels.

Pharmacological implications follow from the kingdom-specific helicase findings: BRIP1 inhibitors might exert opposing effects on intron G4 retention in Plantae versus promoter G4 in Protozoa, a prediction directly testable *in vivo*.

## Methods

### Genome assembly and annotation

Reference genomes and annotations for 198 species spanning the four eukaryotic kingdoms (Animalia *n* = 86, Fungi *n* = 49, Plantae *n* = 33, Protozoa *n* = 30) were obtained from NCBI Assembly, EnsemblGenomes (Metazoa, Plants, Fungi, Protists), WormBase ParaSite, and the Ginkgo Genome Database (Gu et al., 2022). Species selection was guided by phylogenetic representativeness within each kingdom, prioritising chromosome-level assemblies with curated gene annotations.

### Region extraction

For each species, six genomic regions were extracted from the canonical mRNA isoforms: 5*^′^*UTR, CDS, intron, 3*^′^*UTR, promoter (*−*1000 to *−*1 bp relative to the transcription start site), and intergenic. Region BED files were generated using a custom Python pipeline (script 22) that resolves isoform overlaps by union and filters out features lacking gene-model Parent attributes.

### G4 motif detection and stability classification

G4 motifs were detected with a permissive regex pattern G{2,3}[ACGTN]{1,15}G{2,3}[ACGTN]{1,15}G{2,3}[ACG (allowing G-doublets and G-triplets with loop lengths of 1–15 bp, matching the classifier used in our companion viral G4 analysis; Tanigawa and Iwaki, 2026), followed by G4Hunter scoring in a 25-bp sliding window with a per-base mean threshold *|S| ≥* 0.5 (Bedrat et al., 2016); regex candidates overlapping a G4Hunter window by at least 10 bp on the same strand were retained as “consensus candidates.” Predicted thermodynamic stability was computed by Mergny–Lacroix nearest-neighbour analysis (Mergny and Lacroix, 2003), using the per-tetrad increment of *−*2.23 kcal/mol and a long-loop destabilisation penalty of 1.5 *×* max(0*, L*_total_ *−* 9)*/*3 kcal/mol (where *L*_total_ is the summed loop length across the three inter-tract gaps). Consensus candidates with Δ*G ≤ −*5 kcal/mol were classified as “consensus stable” and used for all downstream analyses; under this filter chain the effective final motif set is dominated by canonical G4 with three G-tetrads and total loop length ≲ 12 bp, and no downstream analysis in this paper counts G-doublets or long-loop candidates in isolation.

### Kingdom-level sign-consistency test for the G4 paradox

For each of the six genomic regions (5*^′^*UTR, CDS, intron, 3*^′^*UTR, promoter, intergenic), the per-species log_2_ IRR of consensus stable G4 (region density divided by whole-genome background) was aggregated to a single kingdom summary by taking the within-kingdom median. A kingdom summary was assigned sign “+” if its median *>* 0.005 log_2_ IRR, “*−*” if *< −*0.005, and “0” otherwise (the 0.005 threshold is a numerical tolerance for exact zero medians produced when *≥* 50% of species in a kingdom have zero stable G4 in the region). Sign-consistency across the four kingdoms was then tested under two rules, both explicitly documented in the reproducibility scripts and deposited alongside the manuscript at the Zenodo archive (DOI 10.5281/zenodo.20336866). *Rule B (strict).* A region is sign-consistent only if all four kingdom summaries are non-zero and share the same sign. *Rule A (tie-permissive).* A region is sign-consistent if all kingdom summaries with non-zero sign share the same sign *and* at least three of the four kingdom summaries are non-zero. Under an independent-signs null with equal probability *±* per kingdom, the per-region one-sided probability of a specific same-sign quartet is (1*/*2)^4^ = 0.0625; the joint probability of observing *≥ k* of six regions sign-consistent by chance is computed from the binomial distribution Binom(6, 0.0625). Both rules and both joint probabilities are reported. Zero-stable-G4 species are common in some kingdom-region combinations (Protozoa intron 15/30, Fungi promoter 8/49 with kingdom median exactly zero because of ties around the background level, Protozoa and Fungi 5*^′^*UTR/3*^′^*UTR each *>* 20%), reflecting genuine biological sparsity rather than missing data.

### Helicase orthologue quantification

Protein FASTAs for each species were annotated with eggNOG-mapper-2.1.13 (Cantalapiedra et al., 2021) against eggNOG-5.0.2 (Huerta-Cepas et al., 2019). G4-resolving helicase orthologues were counted by KEGG K-number matches in the eggNOG output: K10901 (BLM/WRN/RECQ), K15255 (PIF1), K15364 (FANCJ/BRIP1), K11271 (RTEL1), K12823 (DDX5), and K17828 (DDX17). PIF1 is registered in KEGG under two orthologue identifiers, K11385 and K15255; the K11385 cluster returned zero eggNOG-mapper hits in all 198 species, whereas K15255 recovered biologically plausible copy-number variation across kingdoms and was used for the analyses reported here. The sum across all six KEGG IDs defines the “total” helicase count.

### Phylogenetic generalised least squares (PGLS)

A 198-tip species tree was constructed from NCBI Taxonomy by extracting the topology relating the v2 sample species; branch lengths were assigned by chronos calibration using Tanigawa and Iwaki (2026) for the deeper eukaryotic nodes (LECA *∼*1.6 Gya, Opisthokonta *∼*1.1 Gya) and standard reference divergence times for within-kingdom nodes. The 51 v2-new species absent from any prior phylogeny were grafted as polytomies attached to their kingdom-level ancestors. Phylogenetic variance-covariance matrices (*V_ij_* = root-to-MRCA depth) were computed for each kingdom independently. PGLS regression *y* = *β*_0_ +*β*_1_*x*+*ɛ* (*ɛ ∼ N* (0*, σ*^2^*V*)) was solved by generalised least squares *β*^^^ = (*X^T^ V ^−^*^1^*X*)*^−^*^1^*X^T^ V ^−^*^1^*y* with significance assessed by the Wald *t*-statistic on *β*^^^_1_. Two phylogenetic robustness analyses were performed: (i) a polytomy-free re-analysis restricted to the 146 v1-original species (scripts/23_pgls_sensitivity_146.py), and (ii) a re-analysis on a topologically independent backbone derived from the Open Tree of Life synthetic tree v15.1 (Hinchliff et al., 2015) (scripts/24a_build_alternative_tree.py and scripts/24_pgls_alternative_tree.py); concordance across the two trees is summarised in Supplementary Table S4.

*d_N_ /d_S_* **at G4 codons (Nei–Gojobori)**

For each kingdom we selected closely related species pairs within the v2 sample (e.g., Animalia: *C. elegans × C. briggsae*). One-to-one orthologue pairs were identified by shared narrowest-level eggNOG ortholog group (largest tax_id). Protein sequences were aligned pairwise with BioPython’s PairwiseAligner under BLOSUM62 (gap open *−*10, gap extend *−*0.5, global mode), and the protein alignments were back-translated to codon alignments using the corresponding CDS sequences. For each codon position the reference-species codon was classified as “G4-overlapping” (any base overlapping a consensus-stable G4 motif) or “control”. The Nei–Gojobori counts of synonymous and non-synonymous differences were computed under the unweighted-paths variant, then converted to evolutionary distances by Jukes–Cantor correction.

### Software availability

All analysis scripts (Python 3.9, dependencies in requirements.txt) are deposited as the manuscript reproducibility package at the Zenodo archive DOI:10.5281/zenodo.20336866.

### Use of generative AI

In accordance with the Cold Spring Harbor Laboratory Press policy on AI tools and large language models, we disclose the following.

Anthropic’s Claude (model: claude-opus-4-7; 1M-context configuration; accessed via the Claude Code CLI in agentic-coding mode between 2026-04-15 and 2026-08-03) was used by the corresponding author (M.T.) to assist with (i) drafting and revising of Python and shell analysis scripts under explicit, version-controlled human review, and (ii) language editing of English prose at the sentence and paragraph level.

OpenAI Codex (ChatGPT 5.6 Sol, accessed between approximately 2026-07-15 and 2026-08-03) was used independently by the second author (T.I.) for cross-document consistency checks, reproducibility review, and identification of potential methodological and reporting issues, with the outputs consolidated into the coauthor audit report deposited as AUDIT_REPORT.md in the Zenodo archive.

Neither tool was used to generate research data or to make autonomous analytical decisions. The authors reviewed the AI-assisted outputs, verified the principal analyses and claims against the underlying deposited results, and take full responsibility for the final work.

Because the coauthor audit identified discrepancies between certain central claims of this manuscript and the deposited code (see the interim-correction notice on page 2 and AUDIT_REPORT.md), the verification described above should be understood as applying only to those items that the interim-correction notice does not flag as pending re-verification. Full author-level verification of the flagged central-analysis items is deferred to a subsequent version following a clean-environment reconstruction.

## Data access

The 198 reference genomes and annotations used in this study are publicly available from their source databases (Supplementary Table S6); accession numbers are listed in data/v2_inventory.csv of the accompanying Zenodo archive. G4 motif BED files, eggNOG annotations, region BED files, heli-case counts, PGLS results, and *d_N_ /d_S_* tables are deposited at Zenodo under DOI:10.5281/zenodo.20336866.

## Acknowledgments

The authors thank the curation teams of NCBI Assembly, WormBase ParaSite, EnsemblGenomes, and the Ginkgo Genome Database for making the underlying reference data publicly available.

## Author contributions

M.T. conceived the study, designed and constructed the analysis pipelines, executed all computational analyses, and drafted the manuscript. T.I. reviewed the code and the manuscript and contributed through critical discussion. Both authors approve the submitted version.

